# PSGL-1 inhibits HIV-1 infection by restricting actin dynamics and sequestering HIV envelope proteins

**DOI:** 10.1101/2020.05.17.100875

**Authors:** Ying Liu, Yutong Song, Siyu Zhang, Min Diao, Shanjin Huang, Sai Li, Xu Tan

## Abstract

PSGL-1 has recently been identified as an HIV restriction factor that inhibits HIV DNA synthesis and more potently, virion infectivity. But the underlying mechanisms of these inhibitions are unknown. Here we show that PSGL-1 directly binds to cellular actin filaments (F-actin) and restricts actin dynamics, which leads to inhibition of HIV DNA synthesis. PSGL-1 is incorporated into nascent virions and restricts actin dynamics in the virions, which partially accounts for the inhibition of virion infectivity. More potently, PSGL-1 inhibits incorporation of Env proteins into nascent virions, leading to loss of envelope spikes on the virions as shown by Cryo-electron microscopy and super-resolution imaging. This loss is associated with a profound defect in viral entry. Mechanistically, PSGL-1 binds gp41 and sequesters gp41 at the plasma membrane, explaining the inhibition of Env incorporation in nascent virions. PSGL-1’s dual anti-HIV mechanisms represent novel strategies of human cells to defend against HIV infection.

## Introduction

HIV-1 infection redirects the cellular machinery towards mass production of new infectious virions, which renders the virus vulnerable to host defense strategies to undermine this production line^1-6^. Currently we only have a very limited understanding of how the human innate immune system targets these vulnerabilities to restrict viral replication. For example, we do not know the functions and mechanisms of actions of the vast majority of the hundreds of potential antiviral genes upregulated by the interferons (interferon stimulated genes, ISGs), the master regulators of antiviral defense^7^. HIV-1 restriction factors, including TRIM5α, APOBEC3, tetherin, SAMHD1 and SERINC3/SERINC5, are a small number of well-studied ISGs that block HIV-1 replication cycles at specific steps^8^. They employ a variety of mechanisms to exert potent restriction on the virus replication. On the other hand, HIV-1 has evolved countermeasures to evade from these blocks, often by inducing the ubiquitination and degradation or relocalization of restriction factors^9^. Given this reoccurring HIV-1-induced degradation of restriction factors, we conducted a genome-wide proteomic profiling of HIV-1 infection in primary human CD4+ T cells and identified a new HIV-1 restriction factor, P-selectin glycoprotein ligand 1 (PSGL-1)^10^. PSGL-1 inhibits HIV-1 reverse transcription and more potently, the infectivity of progeny virions^10^. Moreover, PSGL-1 is induced specifically by interferon γ (IFN-γ) and mediates the bulk of interferon γ’s anti-HIV-1 activity in primary CD4+ T cells^10^. To overcome this restriction, HIV-1 Vpu binds to PSGL-1 and recruits it to the E3 ligase, SCF ^β-TrCP2^ for ubiquitination and subsequent proteasomal degradation^10^. Vpu-deficient HIV-1 has a significantly increased susceptibility to PSGL-1’s infectivity inhibition^10^. Therefore, blocking the interaction between Vpu and PSGL-1 using small molecules could provide a potential therapeutic strategy to quench the infectivity of HIV-1. IFN-γ has largely been considered as an immunomodulatory cytokine and its direct anti-HIV activity is starting to be appreciated^11^. Our work demonstrates that PSGL-1 is a key mediator of IFN-γ’s antiviral activity in human CD4+ T cells, the mechanism of which would therefore be important for the understanding of IFN-γ’s role in the defense against HIV-1 infection.

PSGL-1 has been known as a leukocyte-specific single-pass transmembrane protein that mediates that adhesion of leukocytes to the endothelium at the inflammation sites^12,13^. This adhesion is dependent on the binding of PSGL-1’s extracellular domain to the cognizant selectin proteins (P-, E- or L-selectins) expressed on the surface of endothelium and the attachment of PSGL-1’s cytoplasmic domain to the cortical actin cytoskeleton via ezrin and moesin proteins ^14,15^. Here we show that the cytoplasmic domain of PSGL-1 can directly bind actin filaments (F-actin) and inhibits F-actin disassembly catalyzed by the actin depolymerization protein cofilin^16^. Consequently, PSGL-1 greatly reduces the actin assembly-disassembly dynamics, which are known be required for HIV-1 DNA synthesis^17^ and the migration of HIV-1 DNA to the nucleus^18^. Remarkably, this restriction of actin dynamics is not only true in the cells, but also in the released HIV-1 virions, which can partially account for the inhibition of virion infection by PSGL-1. Moreover, we observed that PSGL-1 inhibits the incorporation of HIV-1 envelope proteins into nascent virions, which accounts for the majority of the infectivity inhibition. Mechanistically, PSGL-1 binds gp41 protein and sequesters it at the plasma membrane, explaining the defect of envelope incorporation. Overall, PSGL-1’s dual mechanisms of actions represent a novel antiviral defense strategy in the repertoire of ISGs.

### PSGL-1’s modulation of F-actin intensity correlates with its anti-HIV-1 activity

PSGL-1 has been reported to attach to the cortical actin cytoskeleton by interacting with ezrin and moesin proteins that are linked to actin^14,15^. This interaction is dependent on the 66 amino acid-long cytoplasmic domain (CD)^14,15^. We confirmed that PSGL-1 can be co-immunoprecipitated with actin and this co-immunoprecipitation is abolished when the cytoplasmic domain is deleted in **Supplementary Fig. S1a**. Fluorescence staining experiments demonstrated a clear colocalization of PSGL-1 and F-actin, with F-actin visualized either with the fluorescence tag-labelled phalloidin^19^ or with a GFP tagged LifeAct reagent^20^ in **Fig. 1a**. We further observed that in Jurkat cells, overexpression of PSGL-1 led to an enhanced cortical F-actin intensity as shown by imaging (**Fig. 1b-c**) or by flow cytometry (**Fig. 1d and Supplementary Fig. S1b**). Consistent with co-immunoprecipitation results, the cytoplasmic domain of PSGL-1 is responsible for the effect on F-actin in **Fig. 1d and Supplementary Fig. S1b**. CRISPR/Cas9 mediated PSGL-1 knockdown in Jurkat cells led to decreased levels of F-actin in **Supplementary Fig. S1c-d**. In addition, overexpression of PSGL-1 inhibits SDF-1-induced chemotaxis of Jurkat cells^21^ (**Supplementary Fig. S1e**), suggesting that PSGL-1 likely inhibits actin depolymerization and actin treadmilling, a driving force for chemotactic mobility of T cells. This is supported by the phalloidin staining of F-actin, which showed that PSGL-1 enhanced the basic level of F-actin intensity but inhibits the actin dynamics induced by SDF-1 in **Supplementary Fig. S1f**. One minute after SDF-1 treatment, Jurkat cells overexpressing luciferase control showed a rapid increase in F-actin density, consistent with a previous study^18^, but Jurkat cells overexpressing PSGL-1 do not have such an increase (**Supplementary Fig. S1f**).

**Figure 1.**
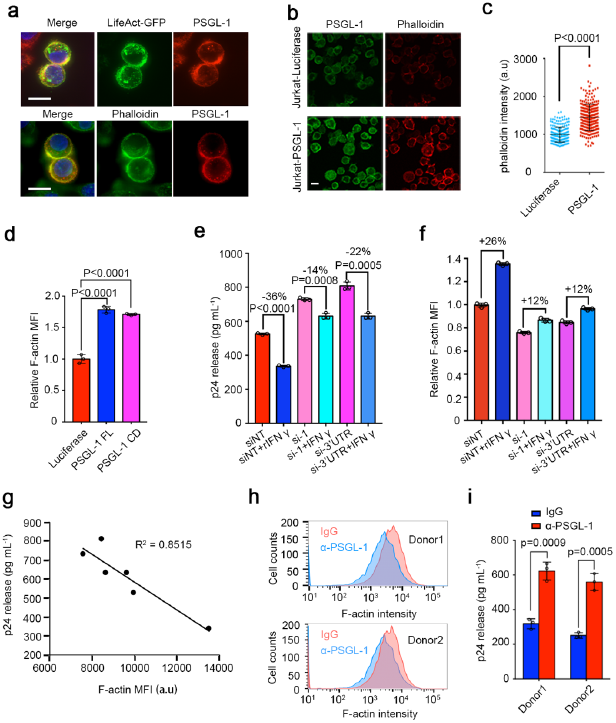
PSGL-1 stabilizes cellular F-actin to restrict HIV infection. **a**, Immunofluorescence staining of PSGL-1 in MAGI cells overexpressing PSGL-1 using anti-PSGL-1 antibody. Upper panel: PSGL-1 (red) colocalizes with LifeAct-GFP (green), which binds F-actin; Lower panel: PSGL-1 (red) colocalizes with phalloidin (green). Scale bar: 10 µm. **b-c**, Immunofluorescence staining of PSGL-1 using anti-PSGL-1 antibody (green) and phalloidin (red) in Jurkat T cells overexpressing luciferase or PSGL-1. Scale bar: 5 µm. The phalloidin intensity of cells in each group were shown in (**c**). **d**, Jurkat T cells overexpressing luciferase, PSGL-1 or PSGL-1 CD (cytoplasmic domain) alone were stained with phalloidin and analyzed with FACS. Relative MFIs were normalized to luciferase group. MFI: Mean Fluorescence intensity. N = 3. **e-f**, Activated primary CD4+ T cells were treated with IFN-γ or mock-treated for 12 h before being electroporated with two different siRNAs targeting PSGL-1 or non-targeting control siRNA (siNT) for 48 h. The cells were then either fixed for phalloidin staining and FACS quantification (**f**) or infected with HIV-1 NL4-3 for 72h before the supernatant being collected for p24 ELISA (**e**). N = 3 for (**e-f**). **g**, Correlation between cellular F-actin intensities and HIV-1 infection rates in (**e-f**). **h-i**, Activated primary CD4+ T cells from two healthy donors were incubated with PSGL-1 antibody or IgG for 2 h before being fixed for phalloidin staining (**h**) or infected with HIV-1 NL4-3 for 72 h before the supernatant being collected for p24 measurement (**i**). N = 3.

We have previously shown that PSGL-1 is induced by IFN-γ and mediates the anti-HIV-1 activity of IFN-γ^10^. Consistently, IFN-γ treatment increased F-actin intensity and this increase is largely abolished when PSGL-1 is knocked down by electroporation of siRNAs (**Fig. 1e-g, Supplementary Fig. S1g**). Importantly, F-actin intensity is negatively correlated with HIV-1 infection, suggesting that PSGL-1’s anti-HIV-1 activity is associated with its modulation of F-actin intensity in **Fig. 1e**. We found that treating primary CD4+ T cells with an antibody that targets the extracellular domain of PSGL-1^22^ can reduce F-actin intensity in the cells, likely due to a conformational coupling between the intracellular and extracellular domains in **Fig. 1h**. This reduced F-actin intensity due to antibody treatment is correlated with an increased HIV-1 infection in **Fig. 1i**, further supporting that PSGL-1 inhibits HIV-1 infection by modulating F-actin intensity.

### PSGL-1 inhibits F-actin depolymerization to restrict HIV-1 reverse transcription

We have previously shown that PSGL-1 expressed in cells infected by HIV (target cells) can inhibit HIV-1 reverse transcription^10^. In addition, previous studies have demonstrated that actin cytoskeleton is required for HIV-1 reverse transcription^17^, providing a potential mechanistic link between PSGL-1’s actin modulation and viral restriction. Indeed, we found that PSGL-1’s inhibition of HIV-1 reverse transcription is dependent on the cytoplasmic domain in **Fig. 2a-b**. Moreover, the cytoplasmic domain itself is sufficient for the inhibition (**Fig. 2b)**. We then performed a series of mutation experiments in the cytoplasmic domain and identified a highly conserved threonine (T393) as a key residue for the binding of PSGL-1 to actin (**Fig. 2c-d, Supplementary Fig. 2**). In contrast, mutations of two nearby conserved serine residues (S385 and S377) had no effect on the binding (**Fig. 2c-d**). Consistently, mutation of T393, but not mutations of S385 or S377, significantly abolished the inhibition of reverse transcription by PSGL-1 (**Fig. 2d**). In addition, PSGL-1 T393A mutant also lost the ability to increase F-actin intensity (**Fig. 2e**). These data support that actin binding and modulation of F-actin intensity by PSGL-1 are required for the reverse transcription inhibition.

**Figure 2.**
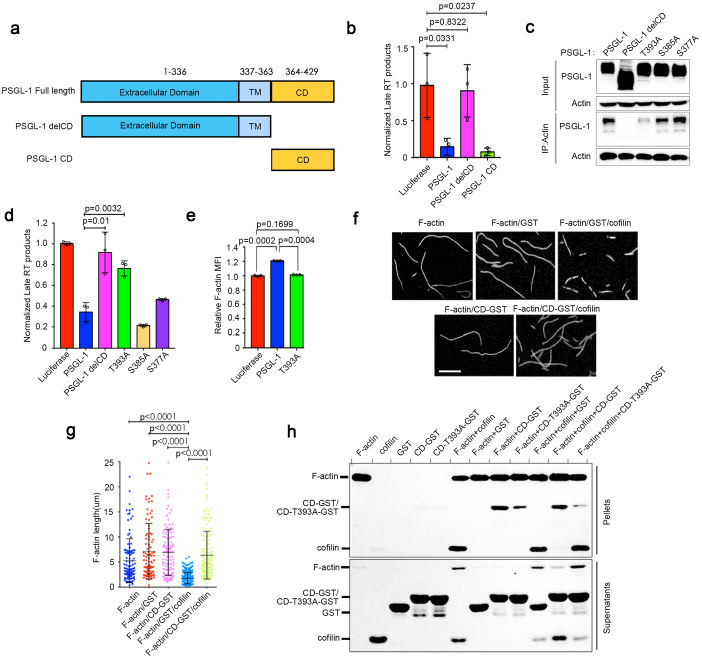
PSGL-1 inhibits F-actin disassembly to inhibit HIV-1 reverse transcription. **a**, Domain structure of PSGL-1, TM: transmembrane domain; CD: cytoplasmic domain; delCD: PSGL-1 with deletion of CD. **b**, MAGI cells overexpressing PSGL-1, PSGL-1 delCD or PSGL-1 CD alone were infected NL4-3 for 12 h to quantitate Late RT product. N=3. **c-d**, MAGI cells transfected with different PSGL-1 constructions for 48h, cells were either immunoprecipitated using anti-actin antibody and protein A agarose beads **(c)** or infected with NL4-3 for 12 h to quantitate Late RT product **(d)**. N=3. **e**, MAGI cells transfected with different PSGL-1 constructs for 48 h then for phalloidin staining and F-actin quantification by FACS. N=3. **f-g**, Purified GST-fusion PSGL-1 cytoplasmic domain or GST alone were mixed with purified and *in vitro* polymerized F-actin for 30 min. The mixtures were later incubated with purified cofilin protein for 1 min and then stopped by phalloidin and measured by fluorescence microscopy **(f)**. Scale bar: 5µm. Quantification of the length of F-actin fibers in **(g)** shows that PSGL-1 CD significantly inhibits the severing of F-actin by cofilin. **h**, Purified GST-fusion PSGL-1 cytoplasmic domain (WT or T393A mutation), purified cofilin and polymerized F-actin were sedimented alone or in the indicated combinations using ultra-centrifugation. The sediments were analyzed by Coomassie staining. The experiments were repeated three times.

Cortical F-actin intensity is determined by the relative speed of actin polymerization and depolymerization reactions in the cells^23^. Since PSGL-1 colocalizes with F-actin, we suspected that it might increase the stability of F-actin by affecting actin depolymerization. We applied a well-established *in vitro* actin depolymerization assay using purified actin filaments and cofilin protein, which is responsible for actin depolymerization^24^. In this assay, we first generated actin filaments and then added purified cofilin to the actin filaments to induce rapid depolymerization of the filaments, which can be observed by fluorescence imaging or by fluorescence TIRF microscopy in real time^24^. We also purified GST-tagged PSGL-1 cytoplasmic domain (CD-GST), which is sufficient to inhibit HIV-1 reverse transcription, due to the difficulty to purify full-length membrane-associated PSGL-1. We found that CD-GST can prevent the depolymerization of F-actin by cofilin. As a control, GST alone had no effect on the depolymerization (**Fig. 2f-g, Supplementary Movies S1-2**). We further utilized an F-actin co-sedimentation assay with purified proteins to show that: (1) CD-GST but not GST alone can be co-sedimented with F-actin; (2) T393A mutation significantly abolished the co-sedimentation of CD-GST with F-actin; (3) CD-GST but not T393A mutant significantly blocked the co-sedimentation of cofilin with F-actin in **Fig. 2h**. These *in vitro* data support that PSGL-1 competes with cofilin for binding to F-actin to block cofilin mediated F-actin depolymerization, leading to an inhibition of HIV-1 reverse transcription.

### PSGL-1 increases F-actin intensity inside HIV-1 virions and affects virion infectivity

We previously showed that PSGL-1 expressed in virus producing cells (producer cells) is packaged into nascent HIV-1 virions and potently restrict the infectivity of the nascent virions even the virions are normalized by p24 levels in **Fig. 3a**^10^. With a β-lactamase assay (BlaM)^25^, we tested cellular entry of the virions packaged with PSGL-1 and found that PSGL-1 significantly inhibited viral entry of the nascent virions (**Fig. 3b-c**). This infectivity inhibition is largely correlated with entry inhibition (**Fig. 3a and 3c**). In addition, both the infectivity inhibition and the entry inhibition are dependent on the cytoplasmic domain of PSGL-1 (**Fig. 3a and 3c**). We reasoned that this might be due to PSGL-1’s effect on F-actin intensity since it has been shown that abundant actin incorporated inside the virions^26^. We confirmed that purified HIV-1 virions contain abundant actin and cofilin in **Supplementary Fig. S3a**. In addition, pre-treatment of the virions with actin polymerization inhibitors, latrunculin A or cytochalasin D promoted viral infection, supporting a role of actin inside the virions in the entry process (**Fig. 3d**). This increased infectivity is associated with increased viral entry as determined using the BlaM assay (**Fig. 3e**). It is worth noting that this promoting effect by actin modulators was not due to actin inhibitors’ effects on the infected cells since treating the cells with the inhibitors at comparable concentrations (the concentration of the compounds in the cell media is much lower than the concentration in the virions due to the dilution of the virions when added to the cells) does not have a promoting effect on the infection (**Supplemental Fig. S3b-c**). Remarkably, the entry and infectivity promotions by these small molecule actin modulators were abolished in PSGL-1-containing virions (**Fig. 3d-e**), suggesting that PSGL-1 has an overriding effect on the actin network in the virions. Super-resolution imaging (STORM)^27^ experiments show that virions packaged with PSGL-1 have significantly higher intensity of F-actin as specifically stained by phalloidin in **Fig. 3f-g**. Importantly, PSGL-1 T393A mutant, which is deficient for actin binding, caused little increase of F-actin intensity, suggesting that this increase is a specific effect associated with actin binding (**Fig. 3f-g**). To verify the STORM imaging results, we applied an ultra-centrifugation method to separate F-actin and actin monomers (G-actin). Western blotting of ultra-centrifugated virions demonstrated increased F-actin levels in virions packaged with PSGL-1, but not those packaged with PSGL-1 T393A mutant (**Fig. 3h**). This loss of ability to promote F-actin intensity of PSGL-1 T393A is associated with a reduction of infectivity inhibition and entry inhibition by this mutant compared to wildtype PSGL-1 (**Fig. 3i-j**), further supporting that PSGL-1 modulation of F-actin intensity inside virions contributes to the inhibition of virion infectivity.

**Figure 3.**
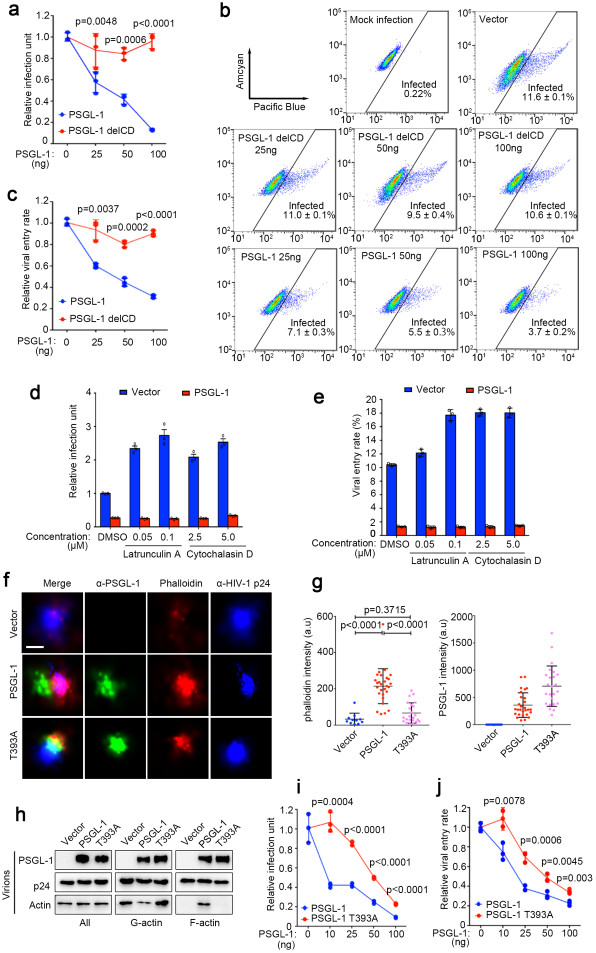
PSGL-1 increases F-actin intensity inside HIV-1 virions and affects virion infectivity. **a**, TZM-bl cells were infected with virions harvested from 293T cells transfected with pNL4-3 and different amounts of plasmids expressing PSGL-1 and PSGL-1 delCD. Empty vector was used to normalize the total transfected DNA. The virions were normalized by p24 ELISA. The infection rates were quantitated with luciferase assay. N=3. **b-c**, Jurkat cells were infected with Vpr-BlaM virions harvested from 293T cells transfected with pNL4-3 and different amounts of plasmids expressing PSGL-1 and PSGL-1 delCD. Empty vector was used to normalize the total transfected DNA. The virions were normalized by p24 ELISA. The infection rates were measured by FACS. N=3. **d**, NL4-3 virions packaged from 293T cells with or without PSGL-1 overexpression were pre-incubated with indicated doses of F-actin inhibitors, latrunculin A or cytochalasin D. TZM-bl cells were infected with treated virions for 48 h before the infection units were measured with luciferase assay. N = 3. **e**, Vpr-BlaM containing virions generated from 293T cells with or without PSGL-1 overexpression were preincubated with latrunculin A or cytochalasin D at indicated concentrations for 2 h. Jurkat cells were infected with the treated virions for 2 h and incubated with β-lactamase substrate overnight before being fixed and analyzed by FACS. N=3. **f**, Virions harvested from producer 293T cells transfected with PSGL-1 or PSGL-1 T393A or an empty vector were pelleted through 20% sucrose cushion, fixed by 4% PFA and stained with antibodies and phalloidin before STORM imaging. Scale bar: 100 nm. **g**, Quantification of the intensities of F-actin or PSGL-1 staining in **(f)** shows that PSGL-1 but not PSGL-1 T393A significantly stabilizes F-actin in virions. **h**, Virions containing PSGL-1 or PSGL-1 T393A were pelleted through 20% sucrose cushion before being sedimented for fractions containing G-actin or F-actin and analyzed by Western blotting. **i**, TZM-bl cells were infected with virions harvested from 293T cells transfected with pNL4-3 and different amounts of plasmids expressing PSGL-1 and PSGL-1 T393A. Empty vector DNA was used to normalize the total DNA transfected. The infection rates were quantitated by luciferase assay. N=3. **j**, Vpr-BlaM containing virions generated from 293T cells transfected with pNL4-3, Vpr-BlaM plasmid and different amounts of PSGL-1 or PSGL-1 T393A plasmids. Empty vector was used to normalize the total DNA transfected. The viruses harvested were normalized by p24 ELISA and used to infect Jurkat cells for 2 h. The cells were then incubated with β-lactamase substrate overnight before being fixed and analyzed by FACS. N=3.

### PSGL-1 inhibits Env incorporation in virions and viral entry

It is worth noting that PSGL-1 T393A, which has little effect on F-actin intensity inside virions, can still strongly inhibit virion infectivity, albeit to a less degree than its wildtype counterpart (**Fig. 3i-j**). This suggests additional, more potent mechanism for the infectivity inhibition by PSGL-1. We found that the virions with PSGL-1 have much less Env proteins, namely gp41 and gp120, than control virions when normalized by p24 levels (**Fig. 4a**), which can explain the entry inhibition we observed earlier (**Fig. 3b-c**). On the other hand, HIV-1 infected Jurkat cells produced virions containing much more Env proteins if PSGL-1 had been knocked out using the CRISPR/Cas9 system (**Fig. 4b-c**). This is true even the virions were normalized by p24, suggesting that PSGL-1’s late effect on Env incorporation is a stronger effect than its early effect. This increase of Env proteins is more significant for Vpu-deficient HIV than the wildtype virus, correlating with Vpu-mediated PSGL-1 degradation (**Fig. 4b-c**). Consistently, Vpu-deficient virions have a more significant increase of infectivity due to PSGL-1 knockout than the wildtype virus (**Fig. 4d**). In fact, we observed a very strong correlation between the gp41 level and virion infectivity in **Fig. 4e**, indicating that the infectivity block by PSGL-1 is due to the loss of Env proteins in the virions. Super-resolution imaging showed that the level of gp41 protein in the virions is significantly increased if the virions were produced from Jurkat cells with PSGL-1 knocked out in **Fig. 4f-g**. On the contrary, if the producer cells overexpress PSGL-1, the gp41 protein levels would be significantly reduced in **Fig. 4h-i**. Gag protein level is also decreased by PSGL-1, but to a less degree than that of gp41 (**Fig. 4h-i**). Consistent with viral infection and entry assays in **Fig. 3a-c**), this effect of PSGL-1 on gp41 level is abolished when CD is deleted (**Fig. 4h-i**), suggesting a requirement for the cytoplasmic domain. We further applied cryo-electron microscopy (Cryo-EM) technique to directly visualize the virions, which showed that PSGL-1-containing virions have much less spikes on the surface of the virions compared to PSGL-1-free virions or PSGL-1 delCD containing virions, supporting an inhibition of Env incorporation in virions by PSGL-1 (**Fig 4j-k**).

**Figure 4.**
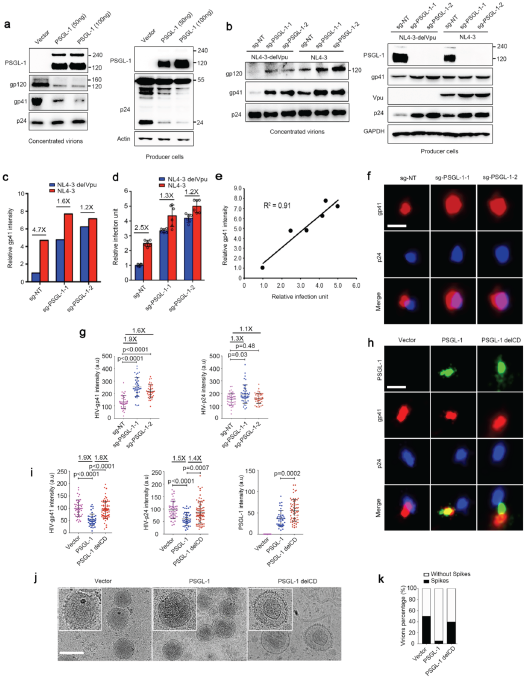
PSGL-1 inhibits Env incorporation in virions and viral entry. **a**, Virions harvested from producer 293T cells transfected with two different doses of PSGL-1 or an empty vector were pelleted through 20% sucrose cushion were analyzed by Western blotting together with the cell lysates. **b**, PSGL-1-knockout Jurkat cells lines with two different sgRNAs or a control cell line with a non-targeting (NT) sgRNA were infected with NL4-3 virus or NL4-3 delVpu virus. The supernatants containing newly released viruses were concentrated and analyzed by Western blotting The cell lysates of the producer cells were also analyzed by Western blotting. **c**, Quantification of the band intensity of gp41 in (**b**) normalized to the intensity of p24. The fold of change between NL4-3 virus and NL4-3 delVpu virus is shown for each cell group. **d**, Virions harvested from infection experiments in (**b**) were normalized by p24 levels measured with ELISA and used to infected TZM-bl cells. The infectivity was measured by luciferase assays. The fold of change between NL4-3 virus and NL4-3 delVpu virus is shown for each cell group. **e**, Correlation analysis between (**c**) and (**d**). **f-g**, Virions from PSGL-1 knockout or control Jurkat cell lines were collected after 5 days post-infection with NL4-3 and pelleted through 20% sucrose cushion were fixed and stained for STORM imaging. Representative images are shown in (**f**). Scale bar: 100 nm. Quantification of STORM images results were showed in (**g**). The ratios between the average values of two groups and the *p* values were shown. **h-i**, Virions from producer 293T cells transfected with PSGL-1, PSGL-1 delCD or an empty vector were pelleted through 20% sucrose cushion were fixed and stained for STORM imaging. Representative images are shown in (**h**). Scale bar: 100 nm. Quantification of STORM images results were showed in (**i**). The ratios between the average values of two groups and the *p* values were shown. **j-k**, Concentrated virions were analyzed by cryo-EM analysis. Representative images were shown in (**j**) and quantification of images of virions shown in (**k**). Scale bar: 100nm.

### PSGL-1 interacts with gp41 and alters cellular localization of gp41

To understand the effect of PSGL-1 on Env incorporation into virions, we first checked if it is due to a defect in the processing of gp160 into gp120 and gp41 using Western blot. The results showed no evidence of such a defect as the ratio of gp120 to gp160 is not altered by PSGL-1 (**Supplementary Fig. S4**). We then tested the interaction between gp41 and PSGL-1 since both are transmembrane proteins. Indeed, immunoprecipitation experiments using either protein as a bait showed that gp41 and PSGL-1 interacts with each other and deletion of CD abolished the interaction, consistent with the infectivity assays (**Fig. 5a**). Moreover, we found two highly conserved leucine residues (L368 and L369) in the cytoplasmic domain (**Supplementary Fig. S2**) to be critical for the interaction between gp41 and PSGL-1 (**Fig. 5a**). Fluorescence staining experiments showed robust colocalization between gp41 and PSGL-1, supporting that the two proteins interact in the cells (**Fig. 5b**). Remarkably, PSGL-1 expression changed the cellular localization of gp41 from mostly intracellular and perinuclear localization to mostly plasma membrane localization (**Fig. 5b-c**). In contrast, PSGl-1 delCD and PSGL-1 LL/AA, both still membrane localized, largely lost the colocalization with gp41 and the ability to relocate gp41 from perinuclear localizations to the plasma membrane. In addition to the CXCR4-tropic NL4-3 strain, PSGL-1’s inhibition of virus entry and effect on gp41 localization also apply to CCR5 strains such as YU2 and NL(AD8) (**Supplementary Fig. S5a-d**). These data suggest a model that PSGL-1 interacts with gp41 in a C-terminal domain-dependent fashion, which sequesters gp41 in the plasma membrane and inhibits its incorporation into nascent virions. Supporting this model, PSGL-1 LL/AA, which cannot bind and relocate gp41, lost the ability to inhibit virion incorporation of Env proteins as shown by Western blotting, super-resolution imaging and Cryo-EM analysis (**Fig. 5d and Supplementary Fig. S5a-d, S6a-d**). Consistently, the infectivity inhibition of PSGL-1 LL/AA is also largely lost due to the mutations (**Fig. 5e and Supplementary Fig. S5a-d**). In comparison, the actin binding and F-actin promoting activity of PSGL-1 LL/AA remain unaffected (**Supplementary Fig. S6e-f**). In contrast to the important role of the LL motif, a mutation previously shown to affect PSGL-1 dimerization (C310A)^28^ or triple mutations previously shown to affect PSGL-1’s co-clustering with Gag (RRK 334/337/338 to AAA or 3A mutations)^29^ do not seem to have an effect on the infectivity inhibition of PSGL-1 (**Supplementary Fig. S7**). How does PSGL-1’s interaction with gp41 excludes Env from being incorporated into nascent virions? A recent study showed that Env is first transported to the plasma membrane and then is endocytosed to the endosomal recycling compartment to assemble with Gag before being released. This trafficking was shown to be required for the viral incorporation of Env^30^. We applied the same Env construct incorporating fluorogen activating peptide (FAP) tags, which allows pulse labeling of Env protein on the cell surface with a membrane impermeable fluorogen. Consistent with the previous report, we observed a rapid internalization of cell surface Env, while this internalization was inhibited by PSGL-1, which strongly co-localizes with Env on the cell membrane (**Fig. 5f**). In contrast, PSGL-1 delCD and PSGL-1 LL/AA are not able to inhibit this internalization. These data together support that a specific interaction between PSGL-1 and gp41 of Env is crucial to the infectivity inhibition by PSGL-1.

**Figure 5.**
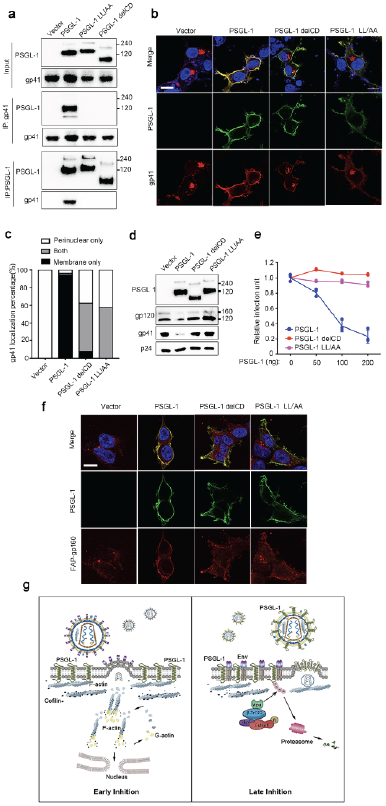
PSGL-1 binds HIV Env and sequesters Env at the plasma membrane. **a**, 293T cells were separately transfected with pNL4-3 proviral plasmid or plasmids expressing PSGL-1, PSGL-1 delCD, PSGL-1 LL/AA or an empty vector. One day after the transfection, PSGL-1 or gp41 were immunoprecipitated with protein A agarose beads with anti-PSGL-1 antibody or anti-gp41 antibody respectively and then the beads were mixed with cell lysates from transfection of the other protein in the IP. The cell lysates and the precipitated proteins were analyzed by Western blotting. **b**, 293T cells transfected with pNL4-3 and PSGL-1 or PSGL-1 delCD or PSGL-1 LL/AA or an empty vector were fixed and stained with anti-gp41 (red), anti-PSGL-1 (green) antibodies and DAPI (blue) and quantification of gp41 cellular localization of each sample was shown in (**c**). Scale bar: 5µm. N=40. **d**, Virions harvested from producer 293T cells transfected with PSGL-1 or PSGL-1 delCD or PSGL-1 LL/AA or an empty vector were pelleted through 20% sucrose cushion were analyzed by Western blotting. **e**, TZM-bl cells were infected with virions harvested from 293T cells transfected with pNL4-3 and different amounts of plasmids expressing PSGL-1, PSGL-1 delCD, PSGL-1 LL/AA. Empty vector was used to normalize the total transfected DNA. The virions were normalized by p24 ELISA. The infection rates were quantitated with luciferase assay. N=3. **f**, Endocytosis of Env (gp160) protein in 293T cells as visualized by pulse labeling of FAP-tag in the Env protein. 293T cells were transfected with plasmids expressing PSGL-1, PSGL-1 delCD, PSGL-1 LL/AA and then stained with the fluorogen MG-11p (red) for 5 min. Cells were fixed after a 20-min chase period and stained with an anti-PSGL-1 (green) antibody and DAPI (blue). Scale bar: 5µm. **g**, Model of the mechanisms of the anti-HIV activity of PSGL-1. In the early stage of HIV life cycle, PSGL-1 restricts cortical actin filament to inhibit HIV reverse transcription. In the late stage of the life cycle, PSGL-1 prevents envelope incorporation into the nascent virions and blocks viral infectivity. PSGL-1 also stabilize F-actin inside of nascent virions to affect their infectivity.

## Discussion

Here we demonstrate that PSGL-1 employs at least two different mechanisms to inhibit HIV infection. For the early inhibition of HIV DNA synthesis, PSGL-1 acts by binding with F-actin and restricting cellular actin dynamics (**Fig. 5g**). A single mutation (T393A) of PSGL-1 that abolishes its F-actin binding also abolished its inhibition of HIV DNA synthesis. Consistently, IFN-γ induces PSGL-1 expression and F-actin intensity, which is abolished by transfection of siRNAs targeting PSGL-1. The strong correlation between our *in vitro* actin depolymerization, binding assay and cellular infection experiments supports this actin restriction mechanism. Actin network has been well known be involved in HIV DNA synthesis^17^ and nuclear migration^18^, consistent with PSGL-1’s restriction of these early processes by modulation of actin dynamics. Remarkably, this restriction also extends to actin in the virions. Actin has been reported as one of the most abundant proteins in HIV virions^26^. In addition, various actin interacting proteins such as ezrin, moesin, coronin-1A and cofilin are all known to be present in the virions^26^. However, the role of the actin cytoskeleton network in the virions has been elusive. We showed that one can promote viral infectivity by disrupting F-actin network with small molecules, while this promotion is abolished by PSGL-1 due to its overriding F-actin stabilizing activity. This observation suggests a role of actin cytoskeleton in viral entry, the mechanism of which is worth further investigation. PSGL-1 represents a novel restriction mechanism that targets the requirement of cytoskeleton for viral infection, which might be relevant to restriction of other types of viruses since actin dynamics are involved in the infection and pathogenesis of many different viruses^31^. PSGL-1’s well established function is mediating the adhesion of leukocytes at inflamed epithelium and T cell migration^12,13,32,33^. Recently, PSGL-1 has been identified as a novel immune checkpoint molecule that mediates T cell exhaustion^34^, it would be interesting to study the relationship between this immunoregulatory function and F-actin restriction of PSGL-1.

Compared to the inhibition of DNA synthesis, virion infectivity inhibition is a more potent block of HIV replication by PSGL-1. The bulk of this infectivity inhibition cannot be explained by PSGL-1’s restriction of F-actin inside of virions. We found that PSGL-1 interacts with gp41 and sequesters it at the plasma membrane so that the nascent virions would be deficient of gp41 to form the spikes, which are required for efficient viral entry (**Fig. 5e**). Western blotting, super-resolution imaging and Cryo-EM data all corroborate the significant decrease of Env proteins in the virions due to PSGL-1. Consistently, PSGL-1 significantly decreased the entry process in a BlaM assay. The infectivity inhibition is largely correlated with the entry inhibition. More importantly, we showed that PSGL-1 knockout in Jurkat cells promotes Env proteins incorporation into the nascent virions and enhances viral infectivity. These loss-of-function data ruled out the possibility of artifact due to protein overexpression. Structurally, PSGL-1’s interaction with gp41 and its infectivity inhibition both require its C-terminal domain, particularly the LL motif in the domain, which supports a strong correlation between these two activities. Mechanically, how PSGL-1’s interaction with gp41 excludes Env from being incorporated into nascent virions would require further investigation, which might shed light on our understanding of the process of viral assembly^35^. Our data support that PSGL-1’s sequestration of gp41 at the plasma membrane would prevent further endocytosis and assembly of Env. In conclusion, PSGL-1 utilizes dual mechanisms to restrict HIV infection at early and late stages. Since the late inhibition of virion infectivity is more potent, HIV uses its Vpu protein to antagonize PSGL-1. Potentially anti-HIV drugs could be developed to inhibit Vpu’s antagonism of PSGL-1 and incapacitate the virions. During the preparation and review of this study, two papers reported mechanistic studies on PSGL-1’s anti-HIV functions^36,37^. These studies proposed a model emphasizing the physical role of virion-incorporated PSGL-1 in occluding virions from attachment to the target cells. Since different study systems, cell lines and different protein expression levels were used in the three studies, further investigation is warranted to incorporate and clarify these findings.

## Author Contributions

XT conceived and supervised the project; YL conducted most of the experiments with the help from SZ; YS and SL conducted Cryo-EM analysis; MD and SH contributed reagents and expertise on the in vitro actin experiments. XT and YL wrote the manuscript.

## Acknowledgements

This work was supported by China National Funds for Excellent Young Scientists (31722030) to XT and grants from Beijing Advanced Innovation Center for Structural Biology, Beijing Frontier Research Center for Biological Structure to XT and SL. We thank Feng Zhou for help with the illustration.

## Supplementary Figures

**Figure S1:**
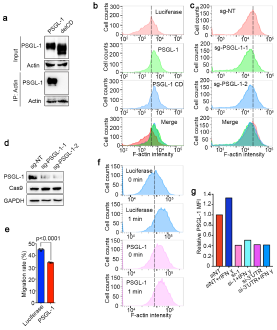
PSGL-1 stabilizes cellular F-actin. **a**, MAGI cells transfected with plasmids expressing PSGL-1 or PSGL-1 delCD for 48 h before the cells were immunoprecipitated with anti-actin antibody and protein A agarose beads. Samples were then analyzed using Western botting. **b**, Jurkat cells overexpressing luciferase, full length PSGL-1 or PSGL-1 cytoplasmic domain (CD) were fixed and stained with phalloidin. The intensity of phalloidin was quantitated with FACS. **c-d**, PSGL-1 knockout Jurkat cells lines were either validated by Western blotting (**d**) or stained with phalloidin for F-actin intensity measurement by FACS (**c**). **e**, Jurkat cells overexpressing luciferase or PSGL-1 were stimulated by SDF-1 for 2 h in a transwell device. **f**, Jurkat cells overexpressing luciferase or PSGL-1 were stimulated with SDF-1 or mock treated for 1 min and immediately fixed and stained with phalloidin for F-actin intensity measurement by FACS. Migrated cells were collected and counted by FACS. **g**, FACS quantification of PSGL-1 expression level of IFN-γ treated and electroporated primary CD4+ T cells in **Fig. 1d and 1e**.

**Figure S2:**
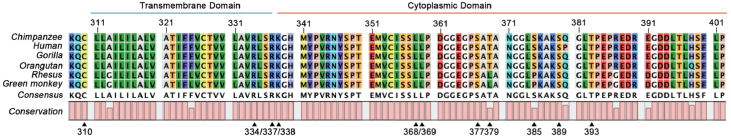
Sequence alignment of PSGL-1. Sequence alignment of the Transmembrane domain and cytoplasmic domain (CD) of PSGL-1 from six primate species. Triangles indicate the residues that were mutated to examine the sequence requirement for actin binding or gp41 binding or mutations that abolished dimerization of gag colocalization.

**Figure S3:**
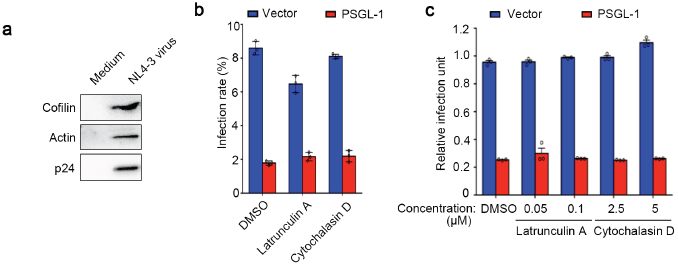
PSGL-1 stabilizes F-actin in HIV virions. **a**, Western blotting analysis shows nascent HIV-1 virions contain abundant cofilin and actin. **b**, Vpr-BlaM containing NL4-3 viruses generated from 283T cells with or without PSGL-1 overexpression were normalized with p24 ELISA. The viruses were used to infect cells treated with latrunculin A or cytochalasin D or DMSO control. The concentrations of the compounds are equal to the higher concentration of each drug in **Fig.3d**. Two hours after infection, the cells were incubated with β-lactamase substrate overnight before being fixed and analyzed by FACs. **c**, TZM-bl cells were treated with actin inhibitors latrunculin A or cytochalasin D or DMSO control at the concentration that is equal to the final concentration of each drug in the cell medium as in **Fig.3e**. NL4-3 viruses generated from 283T cells with or without PSGL-1 overexpression were normalized with p24 ELISA and used to infect the TZM-bl cells pretreated with the indicated compounds. Two days after infection, the infection rate was quantitated using luciferase assay.

**Figure S4:**
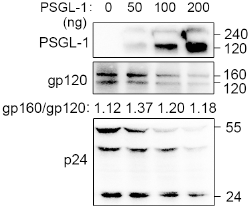
PSGL-1 does not affects Env or Gag processing. 293T cells were transfection with 1 µg pNL4-3 and different amounts of pCMV-PSGL-1 for 48 h before being lysed for Western blotting analysis. Empty vector was used to normalize the total DNA transfect\\

**Supplementary Figure S5:**
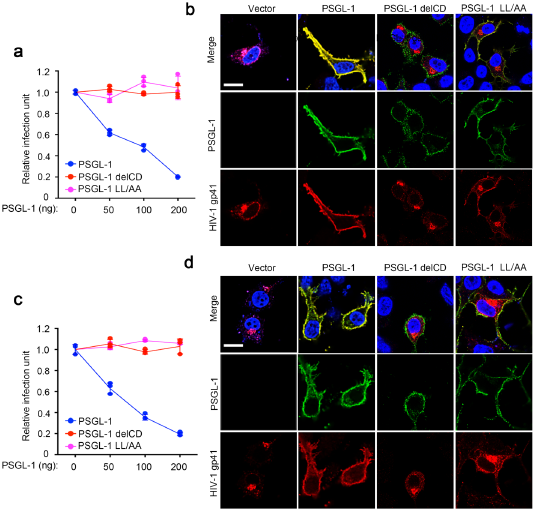
PSGL-1 inhibits R5-tropic virions infectivity and interacts with R5-tropic gp41. TZM-bl cells were infected with virions harvested from 293T cells transfected with two different R5-tropic HIV plasmids pYU2 (**a**) or pNL(AD8) (**c**) and different amounts of plasmids expressing PSGL-1. Empty vector was used to normalize the total transfected DNA. The virions were normalized by p24 ELISA. The infection rates were quantitated with luciferase assays. N=3. 293T cells transfected with pYU2 (**b**) or pNL(AD8) (**d**) and PSGL-1 or an empty vector were fixed and stained with anti-gp41 (red), anti-PSGL-1 (green) antibodies and DAPI (blue). Scale bar: 5µm.

**Figure S6:**
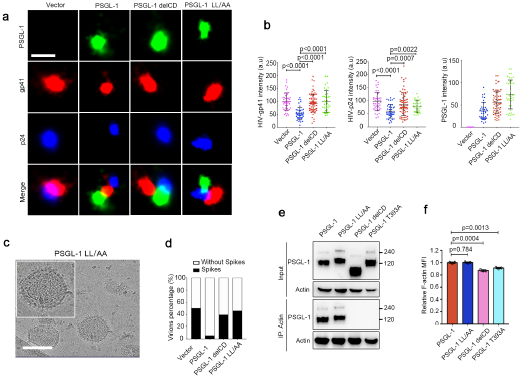
PSGL-1 LL/AA is deficient in gp41 interaction, but not in actin-binding. **a-b**, Virions from producer 293T cells transfected with PSGL-1, PSGL-1 delCD or PSGL-1 LL/AA or an empty vector were pelleted through 20% sucrose cushion were fixed and stained for STORM imaging. Representative images are shown in **(a)** and the quantifications of images are shown in **(b)**. The quantifications were analyzed together with the data in **Fig. 4g** for comparison. **c-d**, Concentrated virions harvested from 293T cells transfected with pNL4-3 and PSGL-1 LL/AA were analyzed by cryo-EM analysis. Scale bar: 100nm. Representative images were shown in **(c)** and quantification of images of virions shown in **(d)**. Scale bar: 100 nm. The quantifications were analyzed together with the data in **Fig. 4i** for comparison. **e**, 293T cells were transfected with plasmids expressing PSGL-1, PSGL-1 delCD, PSGL-1 LL/AA and PSGL-1 T393A. Two days after the transfection, PSGL-1 were immunoprecipitated with protein A agarose beads with anti-Actin antibody. The cell lysates and the precipitated proteins were analyzed by Western blotting. **f**, Jurkat cells transfected with different PSGL-1 constructs for 48 h then for phalloidin staining and F-actin quantification by FACS. N=3.

**Figure S7:**
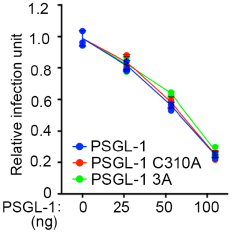
PSGL-1’s dimerization-deficient mutation and gap co-clustering-deficient mutations have no effect on its anti-viral activity. **a**, TZM-bl cells were infected with virions harvested from 293T cells transfected with pNL4-3 plasmids and different amounts of plasmids expressing PSGL-1 or PSGL-1 C336A mutant (dimerization-deficient), PSGL-1 3A mutant (gap co-clustering-deficient). The virions were normalized by p24 ELISA before the infection. The infection rates were quantitated with luciferase assays. N=3.

## Supplementary Movie legends

**Supplementary Movie S1. F-actin+cofilin+GST.avi** and **Movie S2. F-actin+cofilin+CD-GST.avi:**

Purified PSGL-1 cytoplasmic domain in fusion with GST (CD-GST) or GST was mixed with purified and in vitro polymerized F-actin labeled 5-(and-6)-carboxytetramethylrhodamine-succinimidylester for 30 min then the mixtures injected into the flow cell coated with 25 nM N-ethylmaleimidemyosin. Cofilin was injected into the chamber at 10 µM final concentration. Single actin filaments were observed by TIRF illumination with an Olympus IX81 microscope equipped with a 100x oil objective (1.49 NA). Images were collected for 5min with an interval of 3s. The movies were generated with a compression rate of 7 frames/second.

## Methods

### Cells and Cell Culture

293T cells (from ATCC), TZM-bl and MAGI cells (from NIH AIDS Reagents Repository) were cultured in DMEM supplemented with 10% heat-inactivated FBS, 6 mM L-glutamine, and penicillin and streptavidin. Jurkat E6.1 cells were cultured in RPMI-1640 containing 10% FBS, 6 mM L-glutamine and Pen/Strep. Peripheral blood mononuclear cells (PBMCs) were isolated from healthy human donors using Ficoll-Paque PLUS (GE Healthcare). CD4 T cells were purified from PBMCs using anti-human CD4+ magnetic beads (Miltenyi) and then cultured in RPMI supplemented with 10% heat-inactivated FBS, β-mercaptoethanol, and 6mM L-glutamine. Naïve CD4 T Cells were activated with CD3/CD8 beads (Invitrogen) and human recombinant IL-2 (30 U/ml, Roche) for 72 h.

### Plasmids, Transfection and Viral Particle Production

For exogenous expression in mammalian cells, human PSGL-1 or its truncated forms and luciferase were cloned in a pLenti-CMV vector with N-terminal HA tag and FLAG tag, or a PLX-304 vector with a C-terminal V5 tag. The vectors were transfected into cell lines using Neofect transfection reagent following the manufacturer’s protocol (Neofect Biotech, Beijing). For protein purification, human PSGL-1 cytoplasmic domain was cloned in pGEX expression vector with C-terminal GST tag and human cofilin was cloned in pET expression vector with a C-terminal His tag. pCMV-LifeAct-GFP plasmid was purchased from Ibidi. NL4-3 viral particles were produced by transfection of 293T in a T75 flask with 20 µg pNL4-3. SgRNA-expressing and ORF-expressing plasmids were packaged into lentiviral particles in each well of a six-well plate with 10 µg pLKO.1, lentiCRISPR v2 or pLX304 or pCMV backbone vectors, 2 µg pCG-VSV-G, 1 µg pCMV-Tat, 1 µg pCMV-Rev, and 1 µg pCMV-Gagpol. For packaging of Vpr-BlaM viral particles, 10 µg pNL4-3, 3.4 µg pMM310-Vpr-BlaM, and 1.7 µg pAdvantage were transfected into each well of a six-well plate. Twenty-four hours after transfection, the medium was removed. Viral supernatants were collected after another 24 h and filtered at 0.45 µM.

### sgRNA and siRNA

All the siRNAs were purchased from GenePharma, Shanghai. For siRNA nucleofection, the siRNAs were transfected into human CD4+ T cells using the Amaxa Nucleofector following the manufacturer’s protocols (Lonza). The siRNAs used in this study were purchased from GenePharma and the sequences are as below:

siNT sense: UUCUCCGAACGUGUCACGUTT

siNT antisense: ACGUGACACGUUCGGAGAATT

siPSGL-1-1 sense: GCCACUAUCUUCUUCGUGUTT

siPSGL-1-1 antisense: ACACGAAGAAGAUAGUGGCTT

siPSGL-1-3’UTR sense: CAGGAGGCCAUUUACUUGATT

siPSGL-1-3’UTR antisense: UCAAGUAAAUGGCCUCCUGTT

The sgRNA sequences used in this study:

sgNT (Non-targeting control) TTTGAAGTATGCCTCAAGGT

sgPSGL-1-1 TGGGGGAGTAATTACGCACG

sgPSGL-1-2 ATCTAGGTACTCATATTCGG

### Real-time PCR Amplification

For the measurement of viral late RT products and 2-LTR circles, the following primers were used for a TaqMan based qRT-PCR^1^:

Late RT forward primer, 5’-TGTGTGCCCGTCTGTTGTGT-3’

Late RT reverse primer, 5’-GAGTCCTGCGTCGAGAGAGC-3’

Late RT probe, 5’-(FAM)- CAGTGGCGCCCGAACAGGGA-3’

2-LTR forward primer, 5’-AACTAGGGAACCCACTGCTTAAG-3’

2-LTR reverse primer, 5’-TCCACAGATCAAGGATATCTTGTC-3’

2-LTR probe, 5’-(FAM)-ACACTACTTGAAGCACTCAAG-3’

Mitochondrial forward primer, 5′-ACC-CACTCCCTCTTAGCCAATATT-3′

Mitochondrial reverse primer, 5′-GTAGGGCTAGGCCCACCG-3′

Mitochondrial probe, 5′-(TET) CTAGTCTTTGCCGCCTGCGAAGCA (TAMRA)-3′

### Antibodies and Beads

The following antibodies were used for cell cytometry, immunostaining, or Western blotting: anti-PSGL-1 (Santa Cruz Biotech, sc-13535, 1:200), anti-cofilin (Cell Signaling, 5175, 1:500), anti-p24 (NIH AIDS Reagent Program, 1513, 1:1000), anti-Actin (sc-8432, 1:1000), anti-GAPDH (ZSGB-Bio, TA-08, 1:1000), anti-gp41(NIH AIDS Reagent Program, 1242, 1:500), anti-gp120(Abcam, ab21179). For IP and pull-down assay, protein A-agarose beads (Santa Cruz, sc-2001; Bimake, B23202) were used. The fluorescent secondary antibodies used in this study were Alexa Fluor 568 goat anti-human IgG(H+L) (Molecular Probes, A-21090, 1:1000), Alexa Fluor 488 goat anti-mouse IgG(H+L) (Molecular Probes®, A-11001, 1:1000), Alexa Fluor 594 goat anti-mouse IgG(H+L) (Molecular Probes, A-11005, 1:1000), Alexa Fluor 568 goat anti-rabbit IgG(H+L) (Molecular Probes, A-11010, 1:1000), Alexa Fluor 647 goat anti-rabbit IgG(H+L) (Molecular Probes, A-21245, 1:1000), Alexa Fluor 647 rabbit anti-goat IgG(H+L) (Molecular Probes, A-32849, 1:1000). For F-actin staining, Alexa Fluor 568 Phalloidin (Molecular Probes, A-12380, 1:500) was used.

### Immunoprecipitation Assay

For actin IP assay, MAGI cells seeded in 6 cm^2^ plates were transfected with 5 µg pCMV-PSGL-1 or pCMV-PSGL-1 delCD respectively. The cells were lysed with RIPA buffer (50mM Tris-Cl pH7.4, 150mM NaCl, 1% NP-40, 0.5% sodium deoxycholate, 0.1% SDS) plus protease inhibitor cocktail (Bimake, B14001) 48 h after transfection. Cell lysates were incubated with Actin antibody at 4°C overnight. The mixtures were then incubated with 40 µL agarose beads at 4°C for 2 h. The beads were washed three times the next day and boiled in 1X SDS-PAGE buffer at 95 °C for 10 min. For gp41 IP assay. 293T cells in each well of six-well-plates were transfected with 2 µg pNL4-3, pCMV-empty vector, pCMV-PSGL-1, pCMV-PSGL-1 delCD individually. The cells were lysed with binding buffer (50mM Tris-Cl pH7.5, 150mM NaCl, 0.5% NP-40) plus protease inhibitor cocktail at 48 h after transfection. NL4-3-overexpressing 293T cell lysate was mixed with cell lysate of cells expressing PSGL-1, PSGL-1 delCD or pCMV-empty vector and incubated with 40 µl agarose beads and antibodies following the Bimake agarose beads protocol.

### Flow Cytometry

For the SDF-1 stimulation, 2×10^5^ Jurkat cells were stimulated with SDF-1 (400 ng/ml, dissolved in sterilized water) at 37°C. The cells were then fixed quickly using 500µL BD CytoPerm/Cytofix Solution (BD554722) for 20 min at room temperature, washed twice with cold Perm/Wash buffer (BD554723), and stained with Alexa Fluor 568-conjugated phalloidin (Molecular Probes, A-12380) for 30 min at 4°C in the dark. The cells were then washed twice and resuspended in 2% paraformaldehyde in PBS and analyzed on a Becton Dickinson LSR Fortessa flow cytometer. For the antibodies pre-treat experiment, cells were pre-incubated with antibodies at indicated concentrations for 2h at 4°C. Then the cells were fixed and stained with Alexa Fluor 568 Phalloidin.

### Transwell Assay for Chemotaxis

5×10^5^ to 1×10^6^ PSGL-1-overexpressing or Firefly luciferase-expressing Jurkat cells were added to the upper chamber of transwell slide (Costar 3422, Corning), 500 µL RPMI/FBS+20 ng/ml SDF-1 was added to the lower chamber, and cells were allowed to migrate for 2 h at 37 °C in an incubator. Cell numbers were primarily counted with a Bio-Rad TC10 cell counter and then analyzed on a Becton Dickinson LSR Fortessa flow cytometer.

### BlaM Assay

Vpr-BlaM NL4-3 was produced as previously described^2^. For infection of Jurkat cells, 4×10^5^ Jurkat cells in each well of 12-well plates were incubated with NL4-3 virus for 2 h at 37 °C. The cells were washed three times with PBS, and then 0.2 mL 1x CCF4-AM dye solution (Life Technologies, K1095) in phenol-free DMEM/HEPES/2% FBS was added to the cells. The cells were incubated at 11 °C overnight, then washed twice with PBS, fixed in 2% paraformaldehyde in PBS for 20min in dark, and analyzed on a Becton Dickinson LSR Fortessa flow cytometer (Fluorescence channels: Pacific Blue and AmCyan).

### Confocal Microscopy

For F-actin intensity measurement in Jurkat E6.1 cells, 4×10^5^ cells were fixed and stained with anti-PSGL-1 antibody (Santa Cruz Biotech, sc-13535, 1:200) and Alexa Fluor 568 Phalloidin (Molecular Probes, A-12380, 1:500) as previously described. Fixed cells were suspended in 200 µL PBS and dried in a clean cover glass. The cells were analyzed on an ArrayScan VTI 700 (Thermo Scientific, Fluorescence channels: PE and FITC). For F-actin-PSGL-1 colocalization experiment, MAGI cells in 12-well plate were transfected with 1 µg pLifeAct-GFP and 1 µg pCMV-PSGL-1 or pCMV-PSGL-1 alone respectively. Cells were fixed and stained with antibodies. For the gp41-PSGL-1 colocalization experiment, 293T cells in 12-well-plate were transfected with 1 µg pNL4-3 and 1 µg pCMV vector or PSGL-1 constructions. Cells were fixed 20h post infection and stained with PSGL-1 antibody (Santa Cruz Biotech, sc-13535, 1:200), gp41 antibody (NIH AIDS Reagent Program, 1412, 1:500) and imaged as described above. For phalloidin staining, stained cells were imaged using an Olympus FV1200 Confocal Microscope with a 40 NA 0.95 or a 60 NA 1.3 oil objective. For gp41-PSGL-1 colocalization, cells were imaged using Nikon A1R HD25 microscope with a 100 NA 1.3 oil objective. Samples were excited with 480 nm for GFP or 543 nm for Alexa 568 and 594 or 640nm for Alexa 647 and 405 nm for DAPI. F-actin intensities were measured by Imaris software. For the FAP-Env labelling experiment, 293T cells were transfected with FAP-gp160 (kindly provided by Dr. Paul Spearman) and plasmids expressing PSGL-1 constructions at a ratio of 2:1 in 24-well plate. The cells were exposed to 200 nM MG-11p (Spectragenetics) for 5min in dark and then washed with PBS. Then the cells were incubated with DMEM for 20 min at 37 °C and then fixed with PFA and stained with anti-PSGL-1 antibody for imaging analysis.

### STORM Image Acquisition and Analysis

Purified HIV particles were fixed using by BD CytoPerm/Cytofix Solution and adhered to poly-L-lysine solution (SIAMG, P4832) coated glass cover slips for 30 min at room temperature. Cover slips were then blocked using 2% BSA/PBS for 20 min. Particles were stained for PSGL-1 antibody, gp41 antibody and p24 antibody and Alexa Fluor 568 conjugated-anti-human, Alexa Fluor 488 conjugated-anti-mouse and Alexa Fluor 647-anti-goat secondary antibody or phalloidin-Alexa Fluor 568 as indicated. Following immunostaining, particles were washed with PBS and overlaid with image solutions plus 2-mercaptoethanol following Nikon N-STORM manual and imaged using N-STORM microscopy. Images were exported and Fluorescence of each channel were measured by Nikon NIS-Elements AR Analyzer.

### Inhibitor Treatments of Cells

For the SDF-1 treatment, 50ng/mL SDF-1 were added into medium for 1min before Jurkat cells being fixed and stained with phalloidin. For the actin inhibitor treatments of viral particles, supernatants from 293T cells transfected with pNL4-3 with pCMV-vector or pCMV-PSGL-1 were pre-incubated with latrunculin A or DMSO control for 2h at 37 °C or pre-incubated with cytochalasin D at 20 min at 37 °C before being added into cells. The infection rates were determined using luciferase assay or BlaM assay.

### p24 ELISA

For the virus packaged by 293T cells in 12-well plate, cells were transfected with 1µg pNL4-3 proviral plasmids and different doses of plasmids expressing PSGL-1, PSGL-1 delCD or PSGL-1 T393A. Eight hours post transfection, cells were washed twice and the medium were replaced. Two days post transfection, the supernatant was collected for p24 measurement using a commercial p24 ELISA kit (Sinobiological). For the primary CD4+ T cells infection, 1×10^6^ activated CD4+T cells were stimulated by IFN-γ for 24 h before being transfected with 3 siRNAs following Amaxa Nucleofector following the manufacturer’s protocols (Lonza). Two days post transfection, the cells were pre-treated with antibodies for 2 h at 37°C. Then the cells were infected with NL4-3 virus for 6 h, then wash three times with PBS. The supernatant was collected for p24 measurement 72 h post infection.

### Firefly luciferase Assays

Cells transfected with pNL4-3 proviral plasmids and plasmids expressing PSGL-1 or PSGL-1 truncations as described above. Supernatant were collected 48 h after transfection and the virus titrations were measured by p24 ELISA kit. Then TZM-bl cells in 12-well plate were infected with supernatant with equal titration from each group of 293T cells. At 48 h post-infection cells were lysed in cell culture lysis buffer. Virus infectivity was assayed by measuring chemiluminescent firefly luciferase activity using the commercial kit from Promega.

### Protein Purification

The PSGL-1 CD- and cofilin-expressing plasmids were used to transform competent E. coli BL21 strain, which was then grown in LB medium to an OD_600_ of 0.8 and induced with 0.5 mM IPTC at 16 °C overnight. The harvested cells were sonicated in lysis buffer (50 mM Tris HCl pH 7.5, 150 mM NaCl, 2 mM dithiothreitol, and 100nM PMSF). The soluble fraction of cell lysate was purified with the Ni-NTA resin or GST resin (GE) and further fractionated by ion-exchange chromatography (Source Q, GE) and size exclusion chromatography (SD75, GE). Every fraction was collected and validated by SDS-PAGE before the confirmed fractions were combined for later use.

### Fluorescence Microscopy of Actin Filaments

This experiment protocol was as described by Zheng et al^3^. Briefly, 12 µM actin was incubated at room temperature for 30 min in polymerization buffer (10 mM imidazole, 0.2 mM CaCl2, pH 7.0, 2 mM MgCl2, 50 mM KCl, 0.2 mM ATP, 1 mM EGTA, 0.5 mM DTT, and 3 mM NaN3) and with 40 µM GST or PSGL-1 CD-GST or buffer only. The resulting actin filaments (4 µM) were then incubated with 10 µM cofilin for 1 min. An equimolar amount of phalloidin was then added to label actin filaments and stop the reaction. Finally, actin filaments were diluted to 10 nM with fluorescence imaging buffer (10 mM imidazole-HCl, pH 7.0, 50 mM KCl, 1 mM MgCl_2_, 100 mM DTT, 100 mg/mL Glc oxidase, 15 mg/mL Glc, 20 mg/mL catalase, and 0.5% methylcellulose). A sample of 2.5 µL of actin filament mixture was then placed on a poly-L-Lys-coated cover slip and imaged with a 60x oil objective. Image were captured with Image-Pro Express 6.3 software. The length of individual actin filaments was quantified by ImageJ.

### Direct Visualization of Actin Filament Severing and Depolymerization by TIRF Microscopy

This experiment protocol was as described by Zheng et al^3^. Briefly 1.5 µM polymerized actin labeled with 5-(and-6)-carboxytetramethylrhodamine-succinimidylester were incubated with 50 µM GST or PSGL-1 CD-GST for 30 min and then injected into the flow cell coated with 25 nM N-ethylmaleimidemyosin. Cofilin was injected into the chamber at 0.2 µM final concentration. Single actin filaments were observed by TIRF illumination with an Olympus IX81 microscope equipped with a 100x oil objective (1.49 numerical aperture). Images were collected for 5 min with an interval of 1 s using a Photometrics cascade II 512 CCD camera (Major Instruments) with Micromanager software.

### Actin Co-sedimentation Assay

Actin was polymerized as described above. For the F-actin binding complex cotainining 10 µM cofilin, 30 µM GST, CD-GST or CDT393A-GST and 10 µM F-actin filaments in KMEI buffer (10X KEI buffer: 500 mM KCl, 100 mM imidazole pH 7.0, 10 mM MgCl2, 10 mM EGTA) at 30°C for 1 h, and then centrifuged at 10,000g for 30min at 4°C. They supernatant and pellet were separated and boiled for 10 min, followed by electrophoresis and coomassie staining. This experiment was following protocol described by Doolittle et al^4^.

### F-actin isolation

F-actin isolation protocol was followed by Yang et al^5^. Briefly, viruses were filtered thought 0.45µm filters and pelleted through 20% sucrose cushion and then lysed in actin stabilization buffer (100 mM Pipes, pH 6.9, 5% glycerol, 5 mM MgCl2, 5 mM EGTA, 1% Triton X-100, 1 mM ATP and protease inhibitors) pre-warmed at 37°C. Then the lysates were incubated for 10 min at 37°C with shaking and centrifuged at 100000 g for 1 h at 37°C. The pellets containing F-actin were solubilized with actin depolymerization buffer (100 mM Pipes, pH 6.9, 1 mM MgCl2, 10 mM CaCl2 and 5 µM cytochalasin D) and incubated for 1 h on ice. The supernatant (G-actin) and pellet (F-actin) fractions were analyzed by western blotting.

### Cryo-electron microscopy

For virus packaging, supernatants were collected from 293T cells in 15cm^2^ dish transfected as described above before being centrifuged at 4,000 rpm for 5min and then filtered through a 0.45 a 0.45µm filter to remove cell debris. Virion particles was pellet through 20% sucrose cushion and resuspended in 100 µL PBS, stored at 4°C. A 3.5 µL virus sample solution was applied onto a glow discharged copper grid coated with holey carbon (R2/1; Quantifoil, Jena, Germany) prior to plunge-freezing. The grids were blotted for 3.5s, vitrified by plunge-freezing into liquid ethane using a Vitrobot Mark IV (Thermal Fisher Scientific). The grids were imaged using a Tecnai Arctica microscope operated at a voltage of 200 kV equipped with a Falcon III direct electron detector (Thermal Fisher Scientific) at a nominal magnification of 39,000×. Each movie consists of 40 frames was recorded at a calibrated pixel size of 2.69 Å, with 0.05 s/frame exposure, giving a total dose of ∼30e-/Å2 per movie. The electron beam induced motion was corrected by MotionCor2^6^ by averaging 40 frames for each tilt. Image visualization and analysis were performed in IMOD^7^.

### Statistical Analysis

All experiments have repeated at least three times unless otherwise specified. The bar graphs were shown with mean ± SD. Unpaired two-tailed t-test is used to calculate the p-value unless otherwise specified.

